# Multiscale quantification of tissue behavior during amniote embryo axis elongation

**DOI:** 10.1101/053124

**Authors:** Bertrand Bénazéraf, Mathias Beaupeux, Martin Tchernookov, Allison Wallingford, Tasha Salisbury, Amelia Shirtz, Andrew Shirtz, Dave Huss, Olivier Pourquié, Paul François, Rusty Lansford

**Affiliations:** Institut de Génétique et de Biologie Moléculaire et Cellulaire (IGBMC), CNRS (UMR 7104), Inserm U964, Université de Strasbourg, 67400 Illkirch Graffenstaden, France.; Department of Radiology and Developmental Neuroscience Program, Saban Research Institute, Children's Hospital Los Angeles, Los Angeles, CA 90027, USA.; Ernest Rutherford Physics Building, McGill University, 3600 rue University, Montréal, QC, Canada.; Keck School of Medicine, University of Southern California, Los Angeles, CA 90033, USA.; Department of Biological Sciences, University of Southern California, Los Angeles, CA 90089, USA.; Northern Michigan University Computer Science and Mathematics Department, Marquette, MI, 49855, USA.; Department of Genetics, Harvard Medical School and Department of Pathology, Brigham and Woman's Hospital, 77 Avenue Louis Pasteur, Boston, MA 02115, USA.

**Keywords:** confocal microscopy, live imaging, quail embryo, axis elongation, proliferation, PSM, multi-tissue, tissue deformations, morphogenesis

## Abstract

Embryonic axis extension is a complex multi-tissue morphogenetic process responsible for the formation of the posterior part of the amniote body. Cells located in the caudal part of the embryo divide and rearrange to participate in the elongation of the different embryonic tissues (e.g. neural tube, axial and paraxial mesoderm, lateral plate, ectoderm, endoderm). We previously identified the paraxial mesoderm as a crucial player of axis elongation, but how movements and growth are coordinated between the different posterior tissues to drive morphogenesis remain largely unknown. Here we use the quail embryo as a model system to quantify cell behavior and movements in the various tissues of the elongating embryo. We first quantify the tissue-specific contribution to axis elongation by using 3D volumetric techniques, then quantify tissue-specific parameters such as cell density and proliferation at different embryonic stages. To be able to study cell behavior at a multi-tissue scale we used high-resolution 4D imaging of transgenic quail embryos expressing constitutively expressed fluorescent proteins. We developed specific tracking and image analysis techniques to analyze cell motion and compute tissue deformations in 4D. This analysis reveals extensive sliding between tissues during axis extension. Further quantification of “tissue tectonics” showed patterns of rotations, contractions and expansions, which are coherent with the multi-tissue behavior observed previously. Our results confirm the central role of the PSM in axis extension; we propose that the PSM specific cell proliferation and migration programs control the coordination of elongation between tissues during axis extension.

## Introduction

Axis formation in the vertebrate embryo occurs in a head to tail sequence; the head forms first followed by the neck, the thoracic and later on the lumbosacral regions. During this series of events, the different embryonic layers (endoderm, mesoderm and ectoderm) extend toward the posterior pole of the embryo while they progressively organize into more differentiated tissue structures anteriorly. As these structures emerge, they display a stereotypical multi-tissue 3D organization: the neural tube is flanked by two stripes of presomitic paraxial mesoderm (PSM) laterally, by the notochord and endoderm ventrally, and by the ectoderm dorsally.

Technological advances in microscopy have significantly improved our understanding of the morphogenetic events that control vertebrate axis formation, in particular by allowing observation of cellular behaviors over different stages of development. Pioneer studies using frog embryos showed that convergent extension is a central mechanism for the formation of the anterior part of the body axis of vertebrates (Tada and Heisenberg 2012; Shih and Keller 1992a; Shih and Keller 1992b; Bénazéraf and Pourquié 2013). During this process, cells migrate and intercalate, which causes the narrowing of the tissues in one direction and their elongation in the perpendicular direction. In vertebrates, a second phase soon follows during which the axis still extends without considerable change in its width. During this phase, the growth of caudal region of the embryo is thought to be crucial to the elongation process. By deleting caudal structures and developing time-lapse imaging analysis to identify the regions controlling axis elongation in avian embryo, we previously highlighted the critical role of paraxial mesoderm in axis extension and provided evidence of the graded random motility of cells as a primary driver of elongation (Bénazéraf et al. 2010). Although this newly described collective behavior was demonstrated to be important in posterior tissue elongation, it does not explain how movements and growth are coordinated between different tissues in the posterior part of the elongating embryo.

The recent development of tools for analyzing the movements of vast numbers of cells has allowed for better understanding of the complexity of axis morphogenesis. Analyzing cell movements within the tail bud of zebrafish embryos revealed that coherence in collective migration and tissue flow is essential for elongation (Lawton et al. 2013). Interfering with cell/fibronectin interactions disturbed multi-tissue mechanics in this process (Dray et al. 2013). These data suggest large-scale collective migration processes and multi-tissue mechanics are a critical part of the elongation process in fish embryos. We recently established transgenic quails that allow studying global 3D multi-tissue kinetics in real-time in amniote embryos (Huss et al. 2015). These transgenic quails (referred as the H2B-Cherry line) ubiquitously express a nuclear fluorescent protein (H2B-Cherry), which allows the visualization of every nucleus in every tissue of the embryo. Specific algorithms and computational methods have been designed to analyze global tissue deformations in embryos (Blanchard et al. 2009; Rauzi et al. 2015; Rozbicki et al. 2015). However, this type of analyses has not been achieved at the multi-tissue scale in the context of axis elongation of higher vertebrate embryos.

In this work we aim to understand the coordination of growth and movements in the different tissues composing the posterior region of the embryo to allow posterior elongation and organization of the future organs. We use 2P laser imaging on fixed quail embryos to thoroughly assess tissue specific behaviors regarding volume change, cell proliferation and apoptosis at different stages of axis extension. To be able to analyze elongation at the whole structure level dynamically, we use time-lapse imaging of H2B-Cherry transgenic quail embryos to track cell movements and numerically compute related tissue deformations. Our analysis pipeline reveals emergent coordinated motions of multiple tissues, with graded strain rates and highly rotational flows in paraxial mesoderm, and more passive flows in surrounding tissues. Our model indicates that paraxial mesoderm is the driver of extension, as we identified it as the most active zone in terms of cellular flows, cellular proliferation and differential extension, while other adjacent tissues are passively stretched.

## Material and methods

### Quail embryo and embryo culture

Wild-type quail embryos (*Japonica coturnix*) were obtained from commercial sources and from the USC aviary. The *PGK1:H2B-chFP* quail line generation was described previously (Huss et al. 2015) and is maintained in the USC aviary. Embryos were staged according to (Ainsworth et al. 2010; Hamburger and Hamilton 1992) Hamburger and Hamilton, 1951). Embryos were cultured *ex ovo* with filter paper on albumen agar plates according to the EC (Early chick) technique (Chapman et al. 2001).

### Staining and immunodetections

Embryo were collected at the desired stages and fixed overnight in 4% formaldehyde [36% formaldehyde (47608, Sigma) diluted to 4% in PBS]. Blocking and tissues permeabilization were carried out for 2 hours in PBS/0.5% Triton/1% donkey serum. Primary antibodies against Sox2 (1/5000, millipore ab5603) and Bra (1/500, R&D AF2085) were incubated overnight at 4°C. After washing off primary antibody in PBT (PBS/O.1% Triton), embryos were incubated with secondary antibodies (donkey anti goat Alexa 594 and goat anti rabbit Alexa 488, 1/1000, Molecular probes) and DAPI (1/1000, D1306, Molecular probes) overnight at 4°C. The embryos were cleared in U2 scale (Hama et al. 2011) for at least 48h at 4°C and then mounted between slide and coverslip and imaged by confocal/2P microscopy.

### Proliferation and apoptosis analysis

Proliferation was assessed by EdU staining. EdU (Click-iT EdU kit, Cat. #C10083 Life technologies) was diluted in PBS at 10 mM (stock solution). 50 μl of working dilution (500 μM) was dropped on embryos at stage 10. Embryos were incubated for 3, 6, and 9 h, then washed with PBS and fixed in 4% formaldehyde [36% formaldehyde (47608, Sigma) diluted to 4% in PBS]. Apoptosis was assessed using the TUNEL kit (Click-it Plus TUNEL assay (594), Cat. #C106618 Life technologies) after embryo fixation according to manufacturer’s instructions. EdU or TUNEL stained embryos were co-stained with DAPI cleared in U2 scale (Hama et al. 2011)for at least 48h at 4°C and then mounted and imaged by confocal/2P microscopy.

### Imaging

Quail embryos were collected at desired stages (10 to 11HH) with paper filter as described before (Chapman et al. 2001), washed in PBS and either fixed directly in formaldehyde solutions (see above) or cultured on agar albumen plates for EdU incorporation or time-lapse imaging. For fixed tissue imaging embryos were mounted between slide and coverslip, separated by a layer of electrical tape. For live imaging experiments, embryos were cultured at 37C in culture imaging chambers (Lab Tek chambered #1 coverglass slide (Thermo Scientific)) pre-coated with a mix of albumen agar (Chapman et al. 2001). Embryos were imaged using an inverted 780 Zeiss microscope using confocal or two-photon system with 10×/0.45, 20×/0.8 or 25×/0.8 objectives. DAPI (405) and EdU (488) fluorescent signals were separated using 2P online fingerprinting function in ZEN 2011 software (http://zeiss-campus.magnet.fsu.edu/tutorials/spectralimaging/lambdastack/indexflash.html) For time-lapse imaging we imaged several xyz fields of view which were stitched together post-collection. Images were taken every 5-10 um in Z with a time resolution ranging from 5 to 6 minutes.

### Image analysis

Tissue volumes were hand drawn, rendered and volumetric values were calculated using Imaris software (Bitplane). Cell densities were calculated using a variation of k nearest neighbor algorithm (see suppl. methods) based on DAPI stained cell coordinates to segment data-sets using the Spot function of Imaris. The percentage of EdU or TUNEL positive cells was determined by using the Spot function of Imaris. Bra and Sox2 domains were determined using the Surface function of Imaris. New tracking algorithms were designed based on image treatment, nuclei detection, cross-correlation and multi-cellular matching between consecutive frames (see suppl. methods). Tissue movement analysis was computed from cell tracking after flow regularization (using minimization of some energy function), which permits to compute deformation tensors and quantify tissue tectonics (see suppl. methods). Movies were edited using Adobe Premiere, FIJI, and Zeiss ZEN black software.

## Results

### Volume gain varies among tissues during axis extension

As the embryo elongates, the volumes of its different tissues are changing. The relative participation of each tissue in the overall embryonic volume increase is currently unknown. To identify tissue volume changes during axis elongation we imaged a cleared and DAPI stained embryos at different stages of development (11s (11 pairs of somites), 13s and 15s) by 2-photon microscopy. We then analyzed these images by reconstructing the 3D structures of the posterior part of the embryo and manually delineated the different tissue volumes (neural tube, notochord, paraxial mesoderm, progenitor region) based on morphological clues (Fig. 1A). For all tissues, we used the boundary between somite 9 and 10 as an anterior reference because it represents a fixed position easily discernable between embryos. We observed an overall volume increase of the posterior part of the embryo (Fig. 1B). Interestingly, distinct posterior tissues gained volume at different rates (Fig. 1C). For example, the paraxial mesoderm, which constitutes a large part of the whole tissue volume, grows at an average rate of 3.3×10^6^ um^3^/h whereas the neural tube expands at a rate of 1×10^6^ um^3^/h (n=3 embryos for each condition). However, notochord and the axial progenitor zone do not gain significant volume over time.

**Figure 1:**
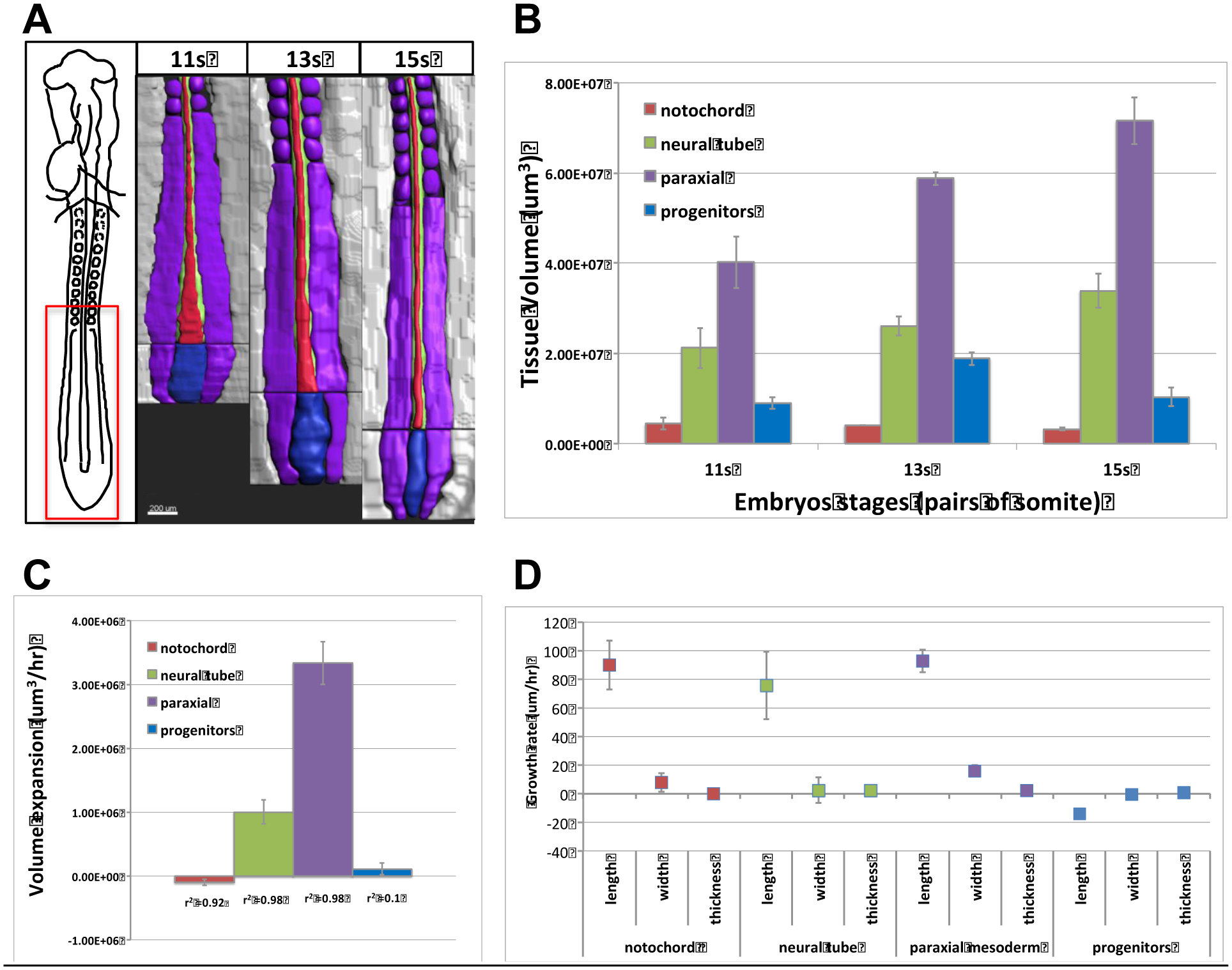
Tissue volume growth during axis extension. A) 3D representation/drawing of different tissues forming the posterior part of quail embryo during axis extension (area of interest is shown in the red rectangle of the scheme on the left, ventral view; anterior side is at the top). The samples are: a representative 11 pairs of somites embryo (left), a 13 pairs of somites embryo (middle), a 15 pairs of somites embryo (right). The progenitor zone is in blue, the paraxial mesoderm is in purple, the notochord is in red and the neural tube is in green. B) Tissue volume measurements for the different stages of development (n=3 embryos per stage). C) Rate of tissue expansion for the different tissues over time (um^3^/hr). r^2^ is calculated for the linear regression of the slope of volume variations. D) Rate of tissue expansion for the different tissue in the different dimensions (*x,y,z*) over time (um/hr). Errors bars are standard errors of the mean.

To characterize if the observed tissue volume variations are anisotropic or isotropic, we measured whether changes are due to gain in length, width or thickness. The analysis shows that the volume expansion is mainly due to an increase in length (Fig. 1D), confirming the strong anisotropy due to anteroposterior extension at the tissue level.

To further improve our volume measurements we used tissue specific marker expression to delimit areas with low morphological cues, for example the boundary between lateral plate and PSM or between the tip of the notochord and progenitor zone. We performed immunostaining of the pan-neural marker Sox2 and pan-mesodermal marker Brachyury to specifically measure the volumes of the neural tube, the notochord, the caudal PSM and the progenitor zone. This analysis confirmed our previous results by showing that the paraxial mesoderm and the neural tube have the fastest volume expansion rate (Sup. Fig. 1). Altogether these results show that individual tissues mostly expand in the anteroposterior direction during axis extension, however, growth rates between tissues differ tremendously.

### Cell density patterns and cell size are tissue specific and conserved across stages

Tissue volume changes can be due to a gain in cell number, or a difference in spacing between cells (Li et al. 2015). To discriminate between these possibilities we calculated the cell density of the posterior part of elongating embryos. Segmentation algorithms were used to localize every nucleus of DAPI-stained and 3D-reconstructed embryo images. Densities were calculated by averaging the distance between 10 neighbor nuclei to generate tissue density maps at different stages of development (Fig. 2A-C). Interestingly, we observed a pattern of regionally conserved densities at different embryonic stages. For example, the neural tube and lateral plate have higher densities than caudal PSM, endoderm and ectoderm. To quantify and compare the tissue specific cell densities we calculated the average density by tissue. The results show that average densities are different between tissues. For instance, the neural tube, notochord and lateral plate have an average densities higher than the paraxial mesoderm, the progenitor zone or the endoderm (Fig. 2D and Sup. Fig. 2). However, when looking at the different stages of development we could not detect significant differences in the average density for each tissue (Fig. 2E and Sup. Fig. 2). Additional to the inter-tissue density differences across stages, our analysis also revealed intra-tissue density differences. For instance, we measured a caudo-rostral gradient of increasing cell density in the PSM at the three different time points (Fig. 2A-C, pink arrowheads). Altogether these results show that cell densities vary within and among tissue types and that these differences are conserved during axis extension. Since our measure of cell densities is based on the measurement of the distance between nuclei we wanted to know if those differences could reflect cell size difference between tissues. To test this possibility we estimated the cell size in different tissues in fixed embryos. Here again this series of measurements show specific distribution of cell size between tissues (Sup. Fig. 3A). For instance endodermal and ectodermal cells are bigger whereas notochord and neural tube contain smaller cells. These differences are conserved in the different stages of elongation (Sup. Fig. 3B). As for the cell density cell size show tissue specific differences that are conserved through the elongation process. Altogether these data strongly suggest that tissue expansion during elongation is not due to cell density or cell size changes and is therefore mostly due to cell number changes.

**Figure 2:**
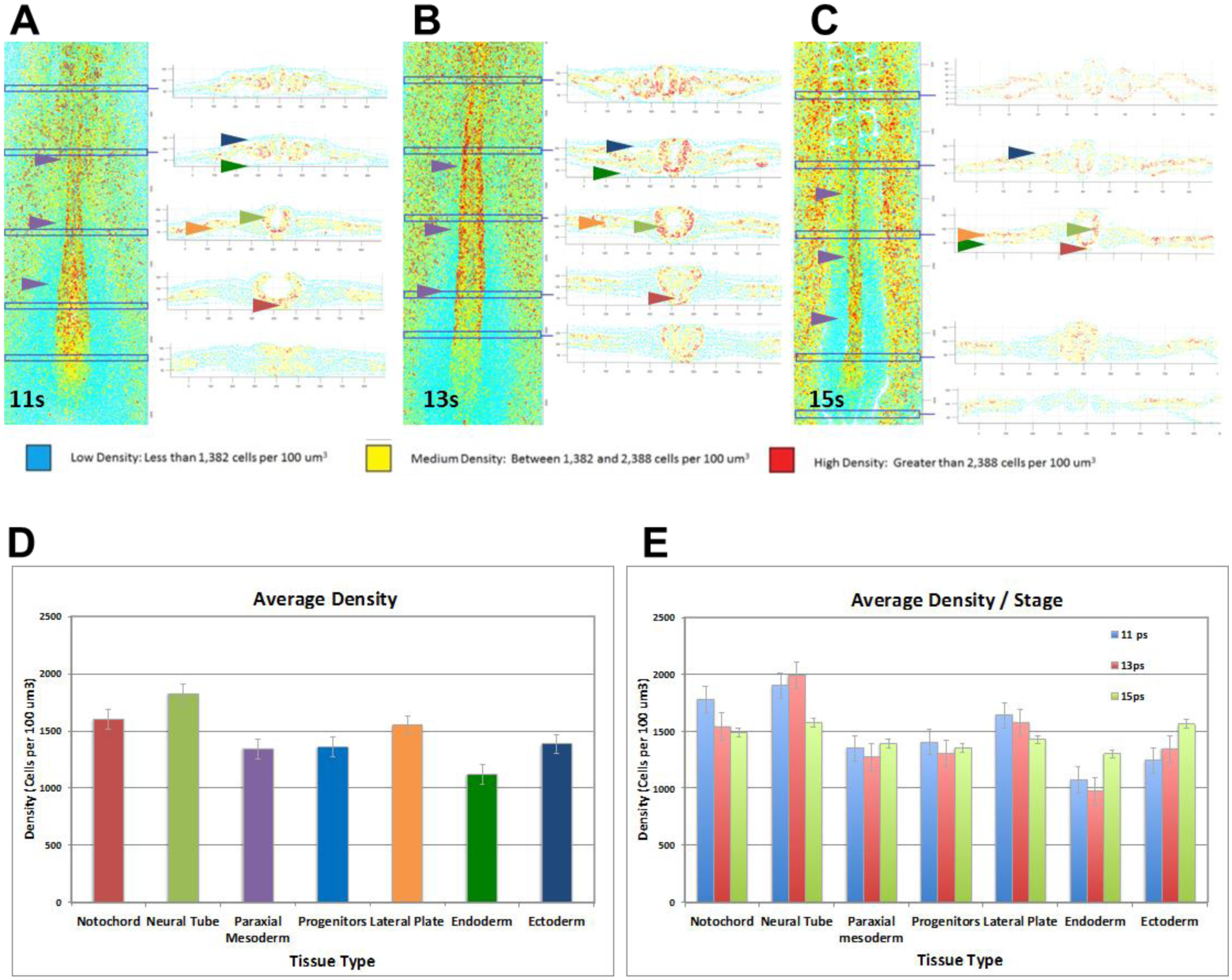
Cell density in the elongating embryo. a-c) 3D density map (dorsal view) and associated transversal sections of representative embryos at different stages 11 pairs of somites (A), 13 pairs of somites (B), 15 pairs of somites (C). Colors represent average densities: cyan codes for density lower than 1382 cells /um3, yellow code for density between 1382 and 2388 cells/um3, red codes for density higher than 2388 cells/um3. The cell densities for PSM, endoderm and ectoderm are low (purple, green, blue arrowheads respectively), cell densities for the notochord, neural tube and lateral plate are medium to high (red, light green and orange arrowhead respectively); note that within the paraxial mesoderm there is a caudal-to-rostral increasing gradient of density (pink arrowheads) D) Average density by tissues. E) Average density per stage (n=3 embryos, Errors bars are standard errors of the mean). Note that we can quantify average tissue density specificities and that these specificities are conserved during the stages analyzed (11,13,15 pairs of somites).

**Figure 3:**
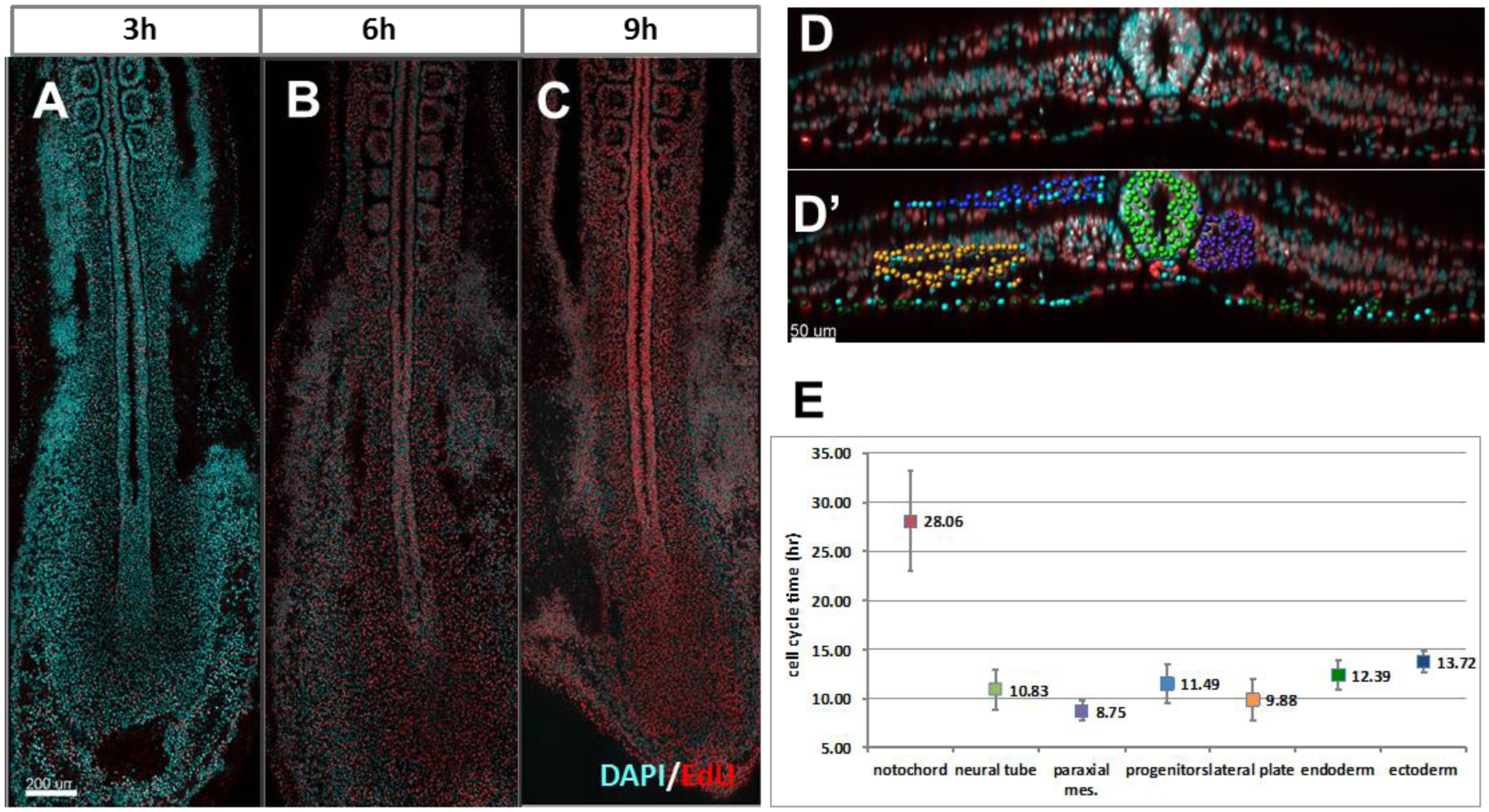
Proliferation analysis during axis extension. A-C) EdU staining (red) and DAPI staining (blue) after different EdU pulse length time on stage 11HH quail embryo: 3h EdU (A), 6 hrs (B), 9hrs (C) (digital confocal coronal slices, anterior is at the top). Note that the longer the pulse the more cells are stained with EdU. D) Example of quantification of 6h EdU positive cells for the different tissues on a digital transversal section of a 3D reconstruction: red is EdU positive, blue is DAPI. D’) EdU positive cells are automatically detected and color-coded depending on the tissue they belong to (paraxial mesoderm is purple, notochord is red, the neural tube is green, ectoderm is dark blue, endoderm is dark green, lateral plate is orange), EdU negative cells are marked by a light blue spot. E) Cell cycle time (in hours) for the different tissues calculated from the rate of percentage of EdU positive cells increase at different pulse times. n=3 to 4 embryos by conditions, errors bars are the standard deviations of the mean.

### Proliferation and apoptosis show tissue specific patterns during elongation

To test whether tissue-specific cell proliferation can be responsible for the different tissue volume growth rates we used cumulative EdU staining and measured tissue cell cycle length (Warren et al. 2009; Nowakowski et al. 1989). We treated embryos for 3h, 6h and 9h with EdU and saw a gradual incorporation of EdU in cells of the posterior part of the embryo (Fig. 3A-C). We then determined the rate of incorporation for each tissue and treatment (Fig. 3D,3D’), which allowed us to calculate the cell cycle times for each tissue. Remarkably, we saw that cell cycle times vary from 8 to 28 h (notochord 28h, ectoderm 13h, endoderm 12h, progenitor zone 11h, neural tube 10h, lateral plate 9h and paraxial mesoderm 8h) (Fig. 3E) When comparing cell cycle times of different tissues to each other we found that, 11 out of 22 comparisons were significantly different (p<0.05, n=3) (Sup. Fig. 4). These results show that in the developing posterior embryo tissues are proliferating at different rates: notochord cells cycle the slowest, endoderm and ectoderm cycle at an intermediate cell cycle time, and cells in paraxial mesoderm, lateral mesoderm, neural tube and axial progenitors zone cycle faster. These results correlate well with the analysis of tissue volume changes since the two tissues that expand the faster, the neural tube and the paraxial mesoderm are also tissue with the shorter cell cycle time (10h and 8h respectively).

**Figure 4:**
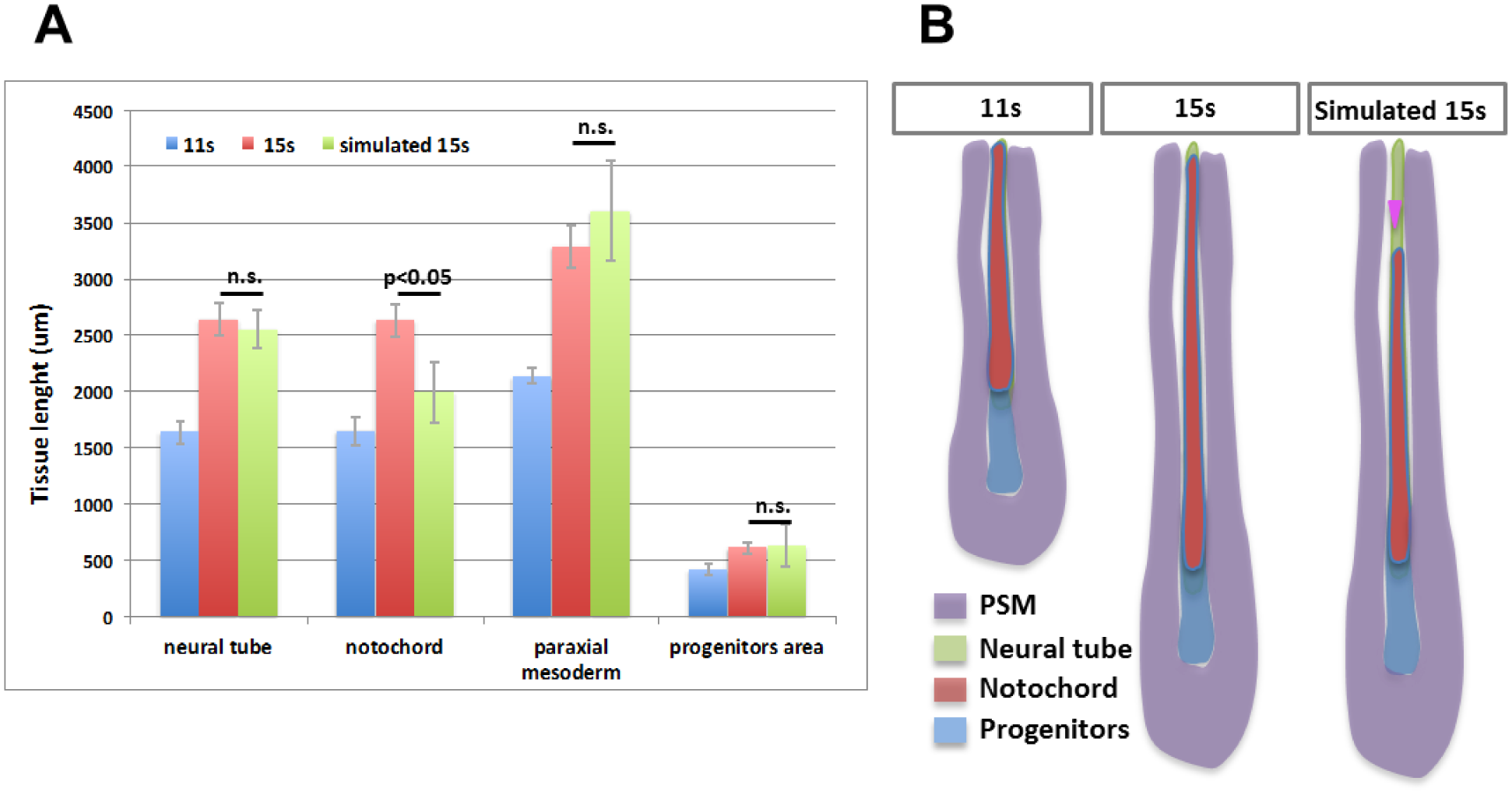
Computational modeling of multi-tissue proliferation and elongation. Comparison of measured and simulated tissue sizes. Average tissue sizes at 11s (blue), 15s (red), simulated by projecting size gain due to tissue specific proliferation (green). The sizes of the PSM, neural tube and progenitor regions are not statistically different between the measurement at 15s and the simulation. By contrast, the size of the notochord is statistically different (p<0.05) between the measurement at 15s and the simulation. B) Schematic representation of the tissue sizes measured and calculated. Note that in the simulation there is a sliding between the notochord and the neural tube and paraxial mesoderm (pink arrowhead).

To see whether there are anteroposterior differences in cell cycle lengths we measured the cell cycle time at the levels of: the anterior PSM (rostral), the caudal PSM (caudal), the progenitor region (very caudal). Although the proliferation speeds within the notochord and endoderm changed significantly along the anteroposterior axis, the tissue cell cycle length did not differ significantly along the A/P axis within the PSM, ectoderm, neural tube or lateral plate mesoderm (Sup. Fig. 4). Altogether our results show that during axis elongation posterior tissues have different cell cycle times.

To assess a potential role of cell death in axis elongation we next examined the rate of apoptosis within the different posterior tissues of the elongating embryo. By combining TUNEL and DAPI staining in 11, 13 and 15s quail embryos we were able to estimate the rate of cell death in 3D (Sup. Fig. 5). The cell death rate did not exceed 5% in any of the tissues and cell death patterns were consistent with published data in chicken embryos (Tenin et al. 2010; Olivera-Martinez et al. 2012). We observed, however, that some tissues exhibited a higher rate of cell death than others. For instance the ectoderm has a cell death rate of 3.6% whereas the axial progenitor zone has the lowest cell death rate of 0.6%. Although cell death only concerns a small proportion of cells, our data shows that cell death seems to be regulated in a tissue specific manner.

**Figure 5:**
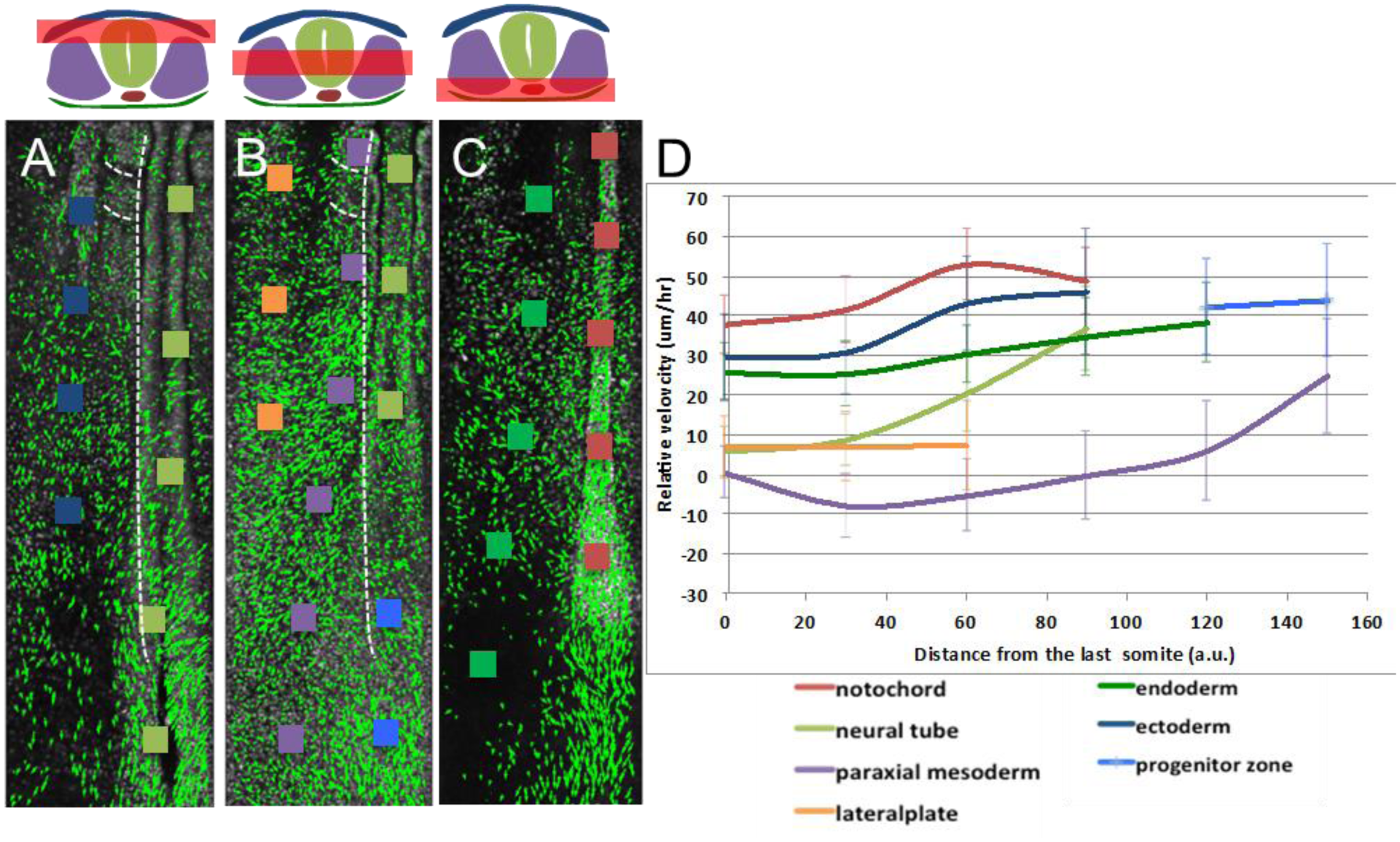
Analysis of tissue movement and tissue sliding during axis extension. A-C: Vector field analysis was computed from *PGK1:H2B-Cherry* transgenic embryo time-lapse data: the green arrows represent the average displacement (length in proportional to distance) and direction of tracked cells. The analysis was performed at different dorsoventral levels (symbolized by the red bar on the schema on the left): dorsal level (A), intermediate level (B) and ventral level (C). Colored squares represents examples of zones in different tissues along the A/P axis in which the analysis has been carried out (purple for paraxial mesoderm, notochord is red, the neural tube is green, ectoderm is dark blue, endoderm is dark green, lateral plate is orange, progenitor region is blue). (D) Averaged differential tissue motilities along the A/P axis. Zones of interest (squares in A-C) were tracked and their motilities were calculated compared to the last formed somite. n=2/3 embryo from the dorsal and ventral side. Errors bars are standard errors of the mean. Distances from the last formed somite to the end of the notochord were normalized.

Altogether our data show that proliferation and cell death display tissue specific pattern in the elongating embryo and therefore these mechanisms can participate in the differential volume growth related to the elongation process. For instance, PSM has a fast proliferation and low cell death rate which can explain the highest volume expansion for this tissue.

### Modeling multi-tissue embryonic growth

After we experimentally characterized different tissue specific cellular behaviors, we wanted to explore their potential influence on axis elongation. To know if the difference in proliferation and expansion can partly explain the multi-tissue kinetics we modeled multi-tissue growth mathematically by fixing the proliferation rate and calculating the elongation rate. We adapted several parameters to facilitate the modeling. We simplified the embryo to a 2D structure composed of four different tissues (paraxial mesoderm, progenitor zone, neural tube and notochord). Since volume expansion is mostly due to growth in the anteroposterior direction (Fig. 1C), we neglected growth in dorsoventral and mediolateral direction. We imposed an elongation rate on each tissue based on experimentally measured proliferation (Fig. 3E), e.g. 28 h for notochord, 11.5h for progenitor zone, 9h for PSM. We then compared the tissue lengths between a virtual 11s embryo grown for 6h (length measured at 11s x rate of proliferation for each tissue) and the experimental measurements of a 15s embryos. The differences between the virtual and experimental measurements of tissue length is therefore indicative of the implication of proliferation in the morphogenetic process of elongation. We find that growth rates of simulated paraxial mesoderm, neural tube and progenitor region are highly similar to the measured growth (p>0.05, n=3) (Fig. 4A). This indicates that proliferation is responsible for a large part of the expansion of these tissues. In contrast, the simulated length of the notochord is significantly shorter than its measured length at 15s and therefore proliferation alone cannot explain the elongation of the notochord (p<0.05, n=3). As the notochord is physically linked to the progenitor region which move toward the caudal direction, our model predict that notochord tissue would slide compare to surrounding tissues such as the paraxial mesoderm (Fig. 4B). Altogether these simulations suggest that differential proliferation is part of a tissue specific coordinative morphogenetic program at work in the posterior part of the developing embryo.

### Embryonic tissues slide during axis elongation

To test the prediction that tissue have differential motions and further understand cell movements and tissue dynamics during axis elongation we used transgenic quail embryos and time-lapse imaging. *PGK1:H2B-ChFP* is a transgenic quail line in which the expression between Histone-2B and mCherry is driven by the ubiquitous pgk promoter. This transgenic model system allows visualization of all cells in every tissue by confocal imaging during early embryogenesis (Huss et al. 2015). Because classic confocal techniques do not allow a deep penetration into the tissue we cultured and imaged embryos from the ventral or dorsal side.

By globally observing tissue kinetics on the ventral side of the embryo we observed that various tissues were elongating at different speed. For example the paraxial mesoderm elongated faster than the notochord (suppl. movie 1), causing a sliding of the two tissues relative to one another as predicted by our theoretical model. Additionally, by using a higher magnification for increased z resolution and by color-coding the z layers we were able to clearly observe differential motion between the endoderm and paraxial mesoderm (suppl. movie 2). In contrast, tissues in the dorsal part of the embryo (neuroectoderm and ectoderm) elongate without any obvious sliding (suppl. movie 3), i.e. at similar speed.

To quantify these diverse tissue kinetics we developed customized cell tracking algorithms to quantitate local differential velocities (detailed in supplemental methods). In each tissue a group of cells located at different positions along the anteroposterior axis (Fig. 5A-C) was tracked, using the last formed somite as a reference point. The last formed somite was chosen as a reference because it is visible from both the ventral and dorsal side and we could thus average tissue movements from embryos filmed from either side. Our analysis shows that all tissues elongate compared to the last formed somite. However, they elongate with different velocity patterns graded along the anteroposterior axis (Fig. 5D). Most tissues (endoderm, ectoderm, lateral plate, notochord) exhibit a weak gradient of velocity (that is, all cells move at the similar velocity), which is suggestive of a cohesive, passive flow. Contrarily, the paraxial mesoderm possesses a very complex behavior with different velocities in its anterior and posterior part. On its anterior part its velocity is slower, different from all other tissues, meaning other tissues dynamically slide pass through it. Local velocity in the anterior gets negative, meaning that anterior PSM cells get closer and closer from the somite and therefore shows tissue compaction. In the posterior part of the PSM we observe a different behavior: there is an increasing gradient of PSM velocity until it catches the velocity of other tissues and therefore move with the other tissues. Altogether our results reveal unexpected tissue choreography of axis elongation with tissues sliding relative to one another. Among these tissues the paraxial mesoderm has very distinct kinetic of movement, consistent with a strong local expansion actively driving elongation.

### Tissue deformations during embryonic axis elongation

Because we found out that axis elongation is defined by differential tissue motions we wanted to examine intra-tissue deformations to see how they could explain this newly described multi-tissue kinetics. We therefore used cell tracking data to compute areas of cell/tissue: 1) rotation 2) compression and expansion, and 3) tensors in the elongating embryo (Fig. 6) (detailed in supplemental methods).

**Figure 6:**
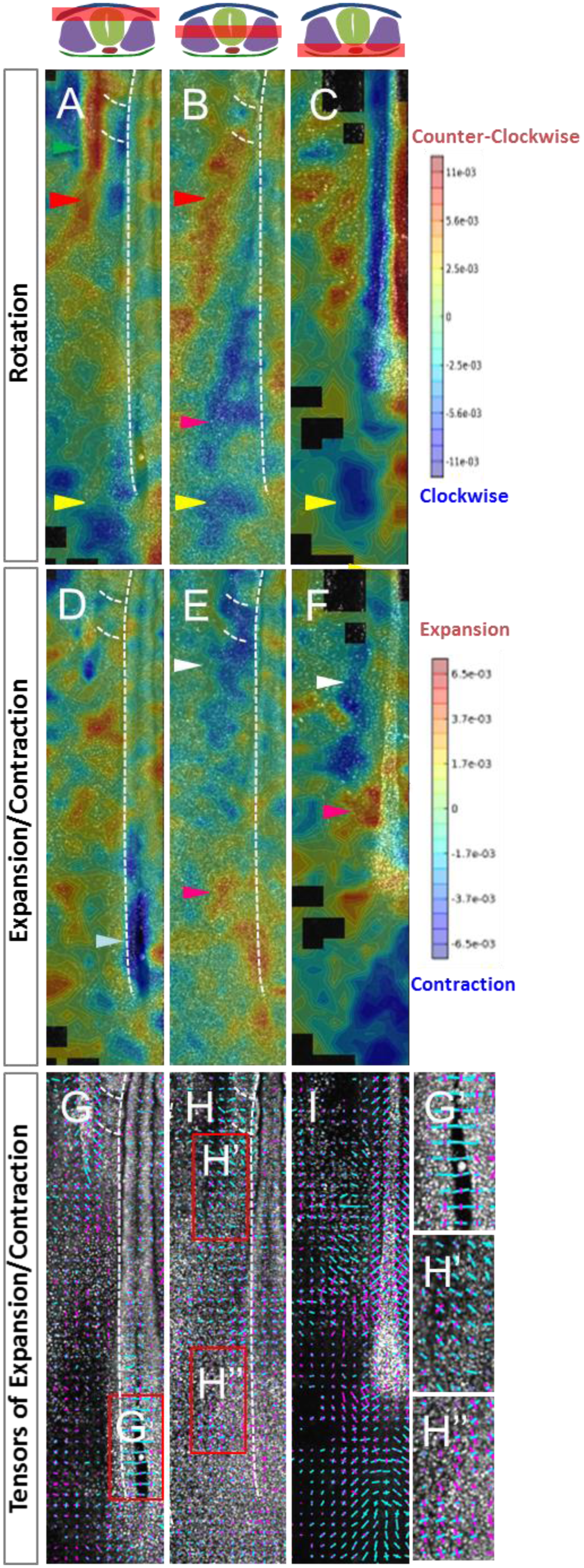
Tissue deformations analysis during axis extension. Rotational analysis (A-C), expansion/contraction (D-F), and tensor maps (G-I) were computed from 3D cell tracking data at different dorsoventral levels from representative embryos: (A,D,G) represents the most ventral level, (B,E,H) intermediate level, (C,F,I) the most dorsal level. In the rotational analysis (A-C) the angle of rotations are color-coded: blue represent the clockwise rotation, red the counter-clockwise rotation. Note that we can see medial to lateral movement in the caudal intermediate part (yellow arrowhead A,B,C) and medial to lateral rotation in the anterior intermediate part (red arrowhead); note also that we also detect sliding at the boundaries between tissues in this analysis (green arrowheads in A show the intermediate mesoderm sliding in between the somites and the lateral mesoderm). In the compaction/expansion map (D-F) contraction and expansion field are color-coded, red representing a expansion and blue a compaction. Note the expansion in the caudal intermediate and ventral level (pink arrowhead), and the contraction in the anterior level, and the neural tube closure (white and light blue arrowheads). In the tensors analysis (G-I) the compaction values are presented in blue and the expansion values in pink, intensities are represented by length, and directionality by their orientation. G’, H’ and H’’ are close up views from G and H. Note the expansion of the caudal ventral region of the embryo has a strong elongation component (A/P axis) (H’’) whereas the contractions in the anterior part of the PSM, or in the caudal neural tube are mainly due to convergence (mediolateral contraction) (G,H’). Black areas represent area with insufficient number of cells to have a statistically significant analysis.

Rotational analysis gives us insights into large-scale cellular flows taking place in the mediolateral axis of the extending embryo. In the posterior part of the analyzed region we observed a general medial to lateral displacement (Fig. 6 A-C, yellow arrowheads). These mediolateral movements are reminiscent of gastrulation movements, as cells leave the progenitor zone to intermix with paraxial mesoderm cells. Interestingly, these movements are not only detected at the level of the mesoderm but also in the most dorsal (ectodermal) and most ventral (endodermal) embryonic levels. In the anterior part of the analyzed region we observed a lateral to medial movement of the mesoderm and ectoderm (Fig. 6 A-B, red arrowheads). These movements resemble convergence, but at the cellular flow level (and not only as local intercalations). Therefore, we see a vortex-type motion that defines elongation at the tissue scale, with cells displaced mediolaterally at the level of the tail of the embryo and lateral to medial at the level of the anterior PSM.

To better understand how differential tissue deformations relates to large-scale tissue movements we created contraction/expansion field maps of the extending embryo (Fig. 6D-F). This analysis shows that most of the dorsal part of the embryo expands with the exception of the closing neural tube, which contracts (Fig. 5D light blue arrowhead). The mesoderm layer (middle layer in the dorsoventral axis) of the embryo behave differently since the posterior paraxial mesoderm and progenitor region expands and the more anterior part of PSM contracts (Fig. 6E,F, pink and white arrowheads respectively).

To determine the directionality of observed deformations we represented the strain tensors (Fig 6 G-I). We observe mediolateral contraction where the neural tube is closing (Fig. 6G,G’). The contraction of the anterior PSM is mainly directed along the mediolateral direction (Fig. 6H,H’) whereas the posterior expansion is directed along the anteroposterior axis (Fig. 6H,H”). These results were confirmed by analyzing different embryos and other time frames of the movies (data not shown and Sup. movies 4).

Interestingly, rotational movements are correlated with expansion and contraction fields. In particular, lateral to medial (counter-clockwise) rotation of the lateral plate and paraxial mesoderm is correlated with compaction in the anterior PSM (Fig. 6B,F red and white arrowheads) whereas mediolateral (clockwise) movements in the mesoderm are correlated with tissue expansion in the caudal PSM and progenitor region (Fig. 6B,F pink arrowheads). This suggests an active role of those rotational motions for tissue elongation.

Altogether our approach allows us to decipher specific intra- and inter-tissue movements in the elongating embryo. Intra-tissue deformations are predominant in the paraxial mesoderm a tissue that has different tissue kinetic than its neighboring tissues (Fig. 5). In the posterior part of the PSM and progenitor zone we observe a clear tissue expansion. In contrast, in the more anterior part of the PSM we observe a convergent cell flow and contractive behavior. These results are consistent with the fact that the PSM deformations could trigger the movements of neighboring tissues.

## Discussion

In this work we used volumetric analysis to show that the posterior elongation in an amniote embryo is the result of complex differential tissues growth. For instance, our results show that the PSM and the neural tube grow more than the neighboring notochord to produce new posterior tissues. The fact that specific cell density is conserved through successive stages suggests that tissue growth is mainly due to the addition of new cells, a fact which is partially confirmed by proliferation data in which we observed that cell cycle time in the PSM and the neural tube are faster than in other tissues such as the notochord. We used H2B-Cherry quail to study cell movements and tissue deformations during axis elongation because this method offers the possibility to dynamically image every cell in all tissues. Our analysis shows that tissues slide along each other during elongation process. In particular differential tissue motion is prominent at the level of the anterior PSM as other tissues slide past the PSM in the caudal direction. Tissue deformation analysis shows that caudal expansion of the PSM tissues is correlated with mediolateral rotational movements and tissue compressions in the anterior PSM tissues that are linked with lateral to medial convergent movements. Altogether our data suggest a model in which the differential expansion of the PSM (due to proliferation and migration/addition of cells in the caudal PSM) induces the coordinated movements between tissues. In the most posterior part of the embryo the anteroposterior expansion of the PSM drives the elongation by stretching the surrounding tissues. In more anterior regions the scenario is different: the PSM tissue contracts and get compacted whereas surrounding tissues still follow the posterior stretch/elongation imposed by the caudal PSM. The difference in behavior between anterior PSM and surrounding tissues is reflected by the tissue sliding that we observed. (Figure 7). This model, that defines caudal PSM role as central in organizing elongation of other tissues, is in line with recent work that outline the presence of regionalized morphogenetic generators in the early developing bird embryo (Loganathan et al. 2014).

**Figure 7:**
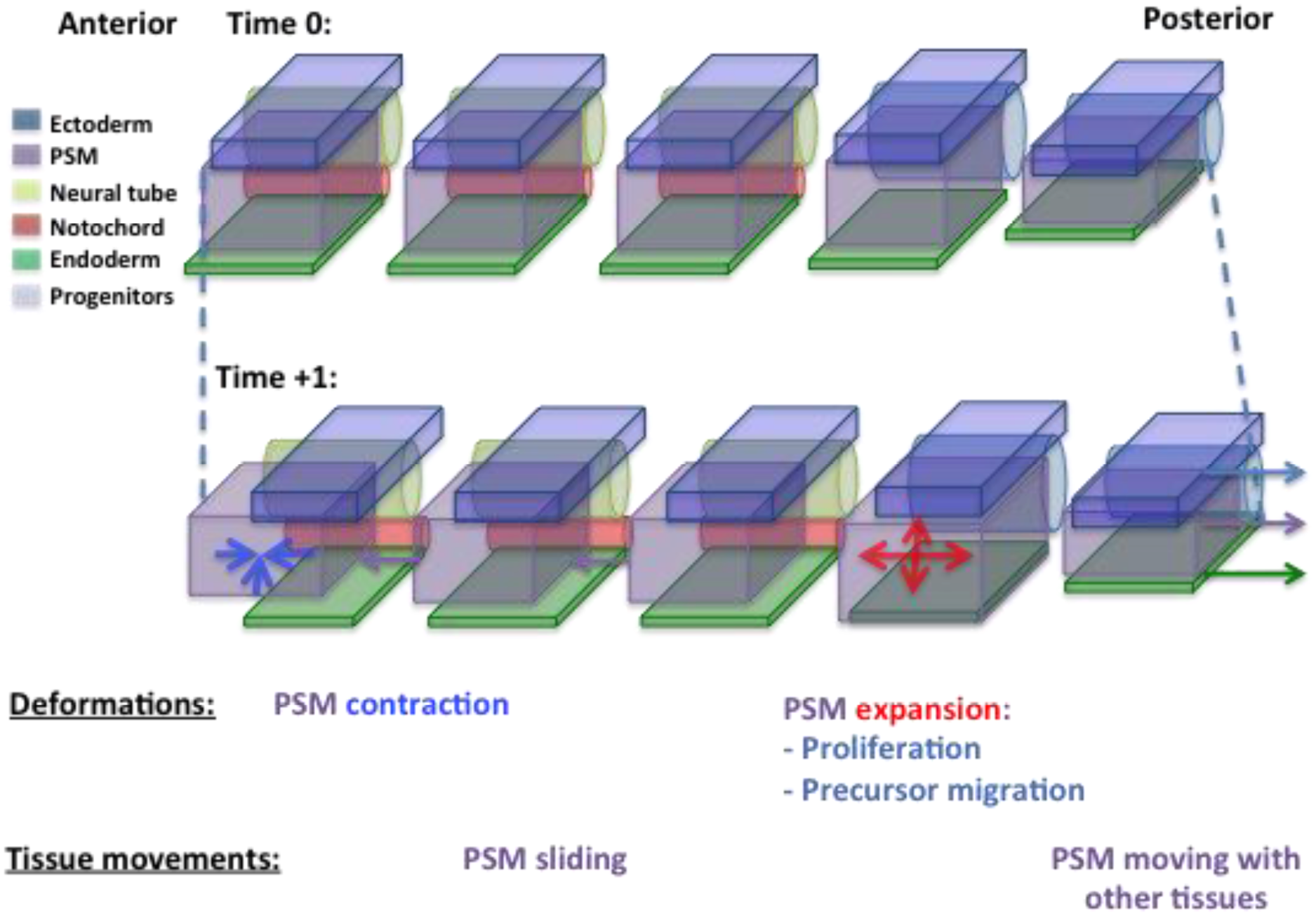
Model of multi-tissue axis elongation. Multi-tissue movements at different anteroposterior levels and time-points showing relative movements of the different tissues (lower part of the panel). Posterior tissues of one side of the embryo are represented from the side view: anterior to the left, posterior to the right (Paraxial mesoderm is purple, notochord is red, the neural tube is green, ectoderm is dark blue, endoderm is dark green). PSM deformations are marked with blue (compression) and red (expansion) arrows. Tissue directional movement is shown by colored arrows: purple for the PSM, blue for ectoderm/progenitors and dark green for endoderm. High proliferation in the PSM and migration in the caudal PSM create local tissue expansion. Compression is taking place in the anterior PSM. The pattern of posterior–expansion/ anterior-contraction create the PSM vortex movement. Posterior PSM expansion cause most caudal tissues to move together because they are physically linked and the rest of PSM to slide with neighboring tissues.

Time-lapse imaging of transgenic avian embryos is an emerging model system to study early morphogenesis events. In particular recent development of transgenic lines has allowed the unraveling of several aspects of early axis formation. Using transgenic chicken ubiquitously expressing membrane-GFP it has been shown that cell intercalation and cell division drive anteroposterior extension of the primitive streak (Firmino et al. 2016; Rozbicki et al. 2015). These studies focus mainly on following cell movements in the epiblast, which is a single cell layered and relatively flat epithelium. To study axis elongation at later stages we needed to use a model system where we could track cells having very different shapes, located in different tissues (epithelial, pseudo-epithelial or mesenchymal) at different dorsal-ventral levels of the developing embryo. We chose to use confocal imaging of transgenic quail expressing H2B-chFP to label the relatively uniform shaped nucleus and specifically designed image treatment that allowed for analysis of cell movement in different tissues. Morphogenetic collective cell movements and tissue deformations are critical aspect of embryonic morphogenesis. Here we demonstrate that the combination of transgenic quail system time-lapse imaging and image analysis allow for studying and quantify movements and deformations at the multi-tissue scale.

Our data on volumetric contribution to axis extension point out that the PSM is a main contributor in quail axis extension. These data are coherent with previous tissue ablation experiments showing that the PSM is central to axis extension in chicken embryo (Bénazéraf et al. 2010). In a recent study it has been shown that zebrafish posterior elongation is mainly due to convergence rather than posterior volumetric growth when compared to mouse embryos (Steventon et al. 2016). This study also point out that the zebrafish PSM region grow much less in comparison to other tissues such as the neural tube, a situation that is very different from the one in bird embryo in which PSM growth is an important part of the overall posterior growth. Therefore axis extension seems to be regulated by different tissue-specific behaviors in different species. The slower growth of zebrafish posterior tissue could be related to the fact that cell cycle in progenitors seems to be slow down in S phase and at the G2/M transition or in G1 phase for the caudal notochord cells (Bouldin et al. 2014; Sugiyama et al. 2014; Sugiyama et al. 2009). Therefore, although zebrafish and quail might have different tissue specific elongation mechanisms the slowing down of the cell cycle in the caudal notochord cells seems evolutionary conserved between these species. We previously shown that global treatment of the chick embryo with cell cycle inhibitors do not alter cellular movements in the PSM (Bénazéraf et al. 2010). However we did not assess for tissue volumetric changes in these experiments. Our present study suggest a crucial role of differential proliferation in tissue kinetics and volumetric growth. Volumetric changes of the different tissues of the posterior part of the embryo in the context of local or global cell cycle inhibition will therefore be critical to functionally assess the role of proliferation in tissue kinetics and axial extension in different species.

Cell mixing in the caudal PSM has been documented extensively (Bénazéraf et al. 2010; Delfini et al. 2005; Kulesa and Fraser 2002; Stern et al. 1988) and is also visible in elongating quail embryo in our study (Sup. Movie 1,2). We proposed that a gradient of random migration is controlling axis extension (Bénazéraf et al. 2010). In the present study, we have quantified tissue deformations and observed that the PSM displays high expansion in its posterior part (where cells move extensively) and contraction in the anterior part (where cell movement diminishes). These deformations are also coherent with the caudo-rostral increasing cell density gradient measured in the PSM. The origins of the forces that trigger compaction have not been identified; they could be coming directly from the cells located anteriorly or/and they can be an indirect effect of long distance forces generated by posterior expansion or the convergence of lateral tissue. The caudal expansion of the paraxial tissue could also result from the integration of different phenomena: (1) the gradient of cell random migration which disperse and reorganizes PSM cells, (2) new cells coming from proliferation (as PSM is actively proliferating in quail) and finally (3) from new cells migrating from the progenitor region (as shown in the rotational analysis). The regulation of the latter has been shown to be an important factor in slowing the elongation process and therefore regulating the size of the body (Denans et al. 2015; Gomez et al. 2008; Iimura and Pourquié 2006). These mediolateral rotational movements that we observed in the caudal part of the embryo are accompanied by lateral to medial movement at the level of the anterior PSM. These movements, also observed in comparable stages in chicken embryo, form large-scale tissue motions as they also involve the ECM (Bénazéraf et al. 2010). Their contributions to posterior axis extension compared to the anteroposterior increasing gradient of tissue expansion is still to be investigated. Interestingly, they could reflect the continuation of existing movements that have been observed during earlier phases of bird development and proposed to be central in early axis extension (Yang et al. 2002; Fleury et al. 2015).

In this study, we show that coordinated and differential tissue movements define amniote axis elongation. Specific pattern of movement and proliferation in the different tissues could create differential expansion and compactions rates leading to sliding between tissue layers. Interestingly the paraxial mesoderm seems to have the most singular behavior compared to other tissues as it contracts anteriorly and expands posteriorly and therefore extending differentially from other tissues. Paraxial mesoderm is covered by ECM matrix such as Fibrillin2 and Fibronectin (Czirók et al. 2004; Martins et al. 2009; Bénazéraf et al. 2010; Rifes and Thorsteinsdóttir 2012). We observed that the Fibronectin movement is more correlated to the PSM cells movements than to other tissues (data not shown). This surrounding layer of ECM might therefore be important in keeping a physical separation between the tissues and in allowing tissue sliding to occur and/or in transmitting forces when tissues are moving together (Araya et al. 2016). In particular, mechanical tissue coupling in the most posterior part of the embryo and mechanical tissue decoupling where tissue slides compared to each other (Fig. 7) could be regulated by different physical properties of the tissues and ECM molecules at the interface between tissues. Data obtained in mouse and zebrafish show that integrin, which are a molecular link between ECM and cells, are required for tissue elongation and tissue mechanics (Girós et al. 2011; Dray et al. 2013). Interestingly, and in coherence with our measures of cell density and tissue deformation in amniotes, biophysical measurements done in zebrafish PSM tissue show that this tissue is stiff anteriorly and more fluid posteriorly (Serwane et al. 2016). This posterior fluidity within the PSM could allow tissue expansion and force generation to deform other tissues. The difference in the nature of the forces created at the interface between the PSM tissue and the matrix or the others tissues at different A/P axis positions might be of importance in generating the tissue kinetics that we observed. Future studies in which we can measure forces exerted between tissues will be of great importance to precisely identify and localize the different factors defining multi-tissue elongation.

## Acknowledgements

We thank Daniela Roellig, Alexis Hubaud, Gonçalo Neto, Ben Steventon and Constance Wu for critically reading the manuscript and members of Rusty Lansford, Scott Fraser, Olivier Pourquié and Paul François teams for suggestions and stimulating discussions during the project.

## Author contributions

Conceived the experiments or provided intellectual input: B.B., R.L, P.F, O.P. Experiments have been performed in the lab of R.L. by: B.B., A.W., T.S., D.H.,. Analyzed the data: B.B., M.B, M.T, T.S, A.S., A.S. Wrote the paper and supervised the work: B.B.

## Bibliography

Ainsworth, S.J., Stanley, R.L. and Evans, D.J.R. 2010. Developmental stages of the Japanese quail. Journal of Anatomy 216(1), pp. 3–15.

Araya, C., Carmona-Fontaine, C. and Clarke, J.D.W. 2016. Extracellular matrix couples the convergence movements of mesoderm and neural plate during the early stages of neurulation. Developmental Dynamics 245(5), pp. 580–589.

Bénazéraf, B., Francois, P., Baker, R.E., Denans, N., Little, C.D. and Pourquié, O. 2010. A random cell motility gradient downstream of FGF controls elongation of an amniote embryo. Nature 466(7303), pp. 248–252.

Bénazéraf, B. and Pourquié, O. 2013. Formation and segmentation of the vertebrate body axis. Annual Review of Cell and Developmental Biology 29, pp. 1–26.

Blanchard, G.B., Kabla, A.J., Schultz, N.L., Butler, L.C., Sanson, B., Gorfinkiel, N., Mahadevan, L. and Adams, R.J. 2009. Tissue tectonics: morphogenetic strain rates, cell shape change and intercalation. Nature Methods 6(6), pp. 458–464.

Bouldin, C.M., Snelson, C.D., Farr, G.H. and Kimelman, D. 2014. Restricted expression of cdc25a in the tailbud is essential for formation of the zebrafish posterior body. Genes & Development 28(4), pp. 384–395.

Chapman, S.C., Collignon, J., Schoenwolf, G.C. and Lumsden, A. 2001. Improved method for chick whole-embryo culture using a filter paper carrier. Developmental Dynamics 220(3), pp. 284–289.

Czirók, A., Rongish, B.J. and Little, C.D. 2004. Extracellular matrix dynamics during vertebrate axis formation. Developmental Biology 268(1), pp. 111–122.

Delfini, M.-C., Dubrulle, J., Malapert, P., Chal, J. and Pourquié, O. 2005. Control of the segmentation process by graded MAPK/ERK activation in the chick embryo. Proceedings of the National Academy of Sciences of the United States of America 102(32), pp. 11343–11348.

Denans, N., Iimura, T. and Pourquié, O. 2015. Hox genes control vertebrate body elongation by collinear Wnt repression. eLife 4.

Dray, N., Lawton, A., Nandi, A., Jülich, D., Emonet, T. and Holley, S.A. 2013. Cell-fibronectin interactions propel vertebrate trunk elongation via tissue mechanics. Current Biology 23(14), pp. 1335–1341.

Firmino, J., Rocancourt, D., Saadaoui, M., Moreau, C. and Gros, J. 2016. Cell Division Drives Epithelial Cell Rearrangements during Gastrulation in Chick. Developmental Cell 36(3), pp. 249–261.

Fleury, V., Chevalier, N.R., Furfaro, F. and Duband, J.-L. 2015. Buckling along boundaries of elastic contrast as a mechanism for early vertebrate morphogenesis. The European Physical Journal. E, Soft Matter 38(2), p. 92.

Girós, A., Grgur, K., Gossler, A. and Costell, M. 2011. α5β1 integrin-mediated adhesion to fibronectin is required for axis elongation and somitogenesis in mice. Plos One 6(7), p. e22002.

Gomez, C., Ozbudak, E.M., Wunderlich, J., Baumann, D., Lewis, J. and Pourquié, O. 2008. Control of segment number in vertebrate embryos. Nature 454(7202), pp. 335–339.

Hama, H., Kurokawa, H., Kawano, H., Ando, R., Shimogori, T., Noda, H., Fukami, K., Sakaue-Sawano, A. and Miyawaki, A. 2011. Scale: a chemical approach for fluorescence imaging and reconstruction of transparent mouse brain. Nature Neuroscience 14(11), pp. 1481–1488.

Hamburger, V. and Hamilton, H.L. 1992. A series of normal stages in the development of the chick embryo. 1951. Developmental Dynamics 195(4), pp. 231–272.

Huss, D., Benazeraf, B., Wallingford, A., Filla, M., Yang, J., Fraser, S.E. and Lansford, R. 2015. A transgenic quail model that enables dynamic imaging of amniote embryogenesis. Development 142(16), pp. 2850–2859.

Iimura, T. and Pourquié, O. 2006. Collinear activation of Hoxb genes during gastrulation is linked to mesoderm cell ingression. Nature 442(7102), pp. 568–571.

Kulesa, P.M. and Fraser, S.E. 2002. Cell dynamics during somite boundary formation revealed by time-lapse analysis. Science (New York) 298(5595), pp. 991–995.

Lawton, A.K., Nandi, A., Stulberg, M.J., Dray, N., Sneddon, M.W., Pontius, W., Emonet, T. and Holley, S.A. 2013. Regulated tissue fluidity steers zebrafish body elongation. Development 140(3), pp. 573–582.

Li, Y., Trivedi, V., Truong, T.V., Koos, D.S., Lansford, R., Chuong, C.-M., Warburton, D., Moats, R.A. and Fraser, S.E. 2015. Dynamic imaging of the growth plate cartilage reveals multiple contributors to skeletal morphogenesis. Nature Communications 6, p. 6798.

Loganathan, R., Little, C.D., Joshi, P., Filla, M.B., Cheuvront, T.J., Lansford, R. and Rongish, B.J. 2014. Identification of emergent motion compartments in the amniote embryo. Organogenesis 10(4), pp. 350–364.

Martins, G.G., Rifes, P., Amândio, R., Rodrigues, G., Palmeirim, I. and Thorsteinsdóttir, S. 2009. Dynamic 3D cell rearrangements guided by a fibronectin matrix underlie somitogenesis. Plos One 4(10), p. e7429.

Nowakowski, R.S., Lewin, S.B. and Miller, M.W. 1989. Bromodeoxyuridine immunohistochemical determination of the lengths of the cell cycle and the DNA-synthetic phase for an anatomically defined population. Journal of Neurocytology 18(3), pp. 311–318.

Olivera-Martinez, I., Harada, H., Halley, P.A. and Storey, K.G. 2012. Loss of FGF-dependent mesoderm identity and rise of endogenous retinoid signalling determine cessation of body axis elongation. PLoS Biology 10(10), p. e1001415.

Rauzi, M., Krzic, U., Saunders, T.E., Krajnc, M., Ziherl, P., Hufnagel, L. and Leptin, M. 2015. Embryo-scale tissue mechanics during Drosophila gastrulation movements. Nature Communications 6, p. 8677.

Rifes, P. and Thorsteinsdóttir, S. 2012. Extracellular matrix assembly and 3D organization during paraxial mesoderm development in the chick embryo. Developmental Biology 368(2), pp. 370–381.

Rozbicki, E., Chuai, M., Karjalainen, A.I., Song, F., Sang, H.M., Martin, R., Knölker, H.-J., MacDonald, M.P. and Weijer, C.J. 2015. Myosin-II-mediated cell shape changes and cell intercalation contribute to primitive streak formation. Nature Cell Biology 17(4), pp. 397–408.

Serwane, F., Mongera, A., Rowghanian, P., Kealhofer, D.A., Lucio, A.A., Hockenbery, Z.M. and Campàs, O. 2016. In vivo quantification of spatially varying mechanical properties in developing tissues. Nature Methods.

Shih, J. and Keller, R. 1992a. Cell motility driving mediolateral intercalation in explants of Xenopus laevis. Development 116(4), pp. 901–914.

Shih, J. and Keller, R. 1992b. Patterns of cell motility in the organizer and dorsal mesoderm of Xenopus laevis. Development 116(4), pp. 915–930.

Stern, C.D., Fraser, S.E., Keynes, R.J. and Primmett, D.R. 1988. A cell lineage analysis of segmentation in the chick embryo. Development 104 Suppl, pp. 231–244.

Steventon, B., Duarte, F., Lagadec, R., Mazan, S., Nicolas, J.-F. and Hirsinger, E. 2016. Species-specific contribution of volumetric growth and tissue convergence to posterior body elongation in vertebrates. Development 143(10), pp. 1732–1741.

Sugiyama, M., Saitou, T., Kurokawa, H., Sakaue-Sawano, A., Imamura, T., Miyawaki, A. and Iimura, T. 2014. Live imaging-based model selection reveals periodic regulation of the stochastic G1/S phase transition in vertebrate axial development. PLoS Computational Biology 10(12), p. e1003957.

Sugiyama, M., Sakaue-Sawano, A., Iimura, T., Fukami, K., Kitaguchi, T., Kawakami, K., Okamoto, H., Higashijima, S. and Miyawaki, A. 2009. Illuminating cell-cycle progression in the developing zebrafish embryo. Proceedings of the National Academy of Sciences of the United States of America 106(49), pp. 20812–20817.

Tada, M. and Heisenberg, C.-P. 2012. Convergent extension: using collective cell migration and cell intercalation to shape embryos. Development 139(21), pp. 3897–3904.

Tenin, G., Wright, D., Ferjentsik, Z., Bone, R., McGrew, M.J. and Maroto, M. 2010. The chick somitogenesis oscillator is arrested before all paraxial mesoderm is segmented into somites. BMC Developmental Biology10, p. 24.

Warren, M., Puskarczyk, K. and Chapman, S.C. 2009. Chick embryo proliferation studies using EdU labeling. Developmental Dynamics 238(4), pp. 944–949.

Yang, X., Dormann, D., Münsterberg, A.E. and Weijer, C.J. 2002. Cell movement patterns during gastrulation in the chick are controlled by positive and negative chemotaxis mediated by FGF4 and FGF8. Developmental Cell 3(3), pp. 425–437.

